# Spinal Cord Injury regulates circular RNA expression in axons

**DOI:** 10.1101/2023.04.26.538466

**Authors:** Mustafa M. Siddiq, Carlos A. Toro, Nicholas P. Johnson, Jens Hansen, Yuguang Xiong, Wilfredo Mellado, Rosa E. Tolentino, Kaitlin Johnson, Gomathi Jayaraman, Zaara Suhail, Lauren Harlow, Kristin G. Beaumont, Robert Sebra, Dianna E. Willis, Christopher P. Cardozo, Ravi Iyengar

**Affiliations:** Pharmacological Sciences & Institute for Systems Biomedicine, Icahn School of Medicine at Mount Sinai, NY, NY; Spinal Cord Damage Research Center, James J. Peters VA Medical Center, Bronx, NY; Department of Medicine, Icahn School of Medicine at Mount Sinai, NY, NY; Burke Neurological Institute, White Plains, NY; Department of Genetics & Genomic Studies, Icahn School of Medicine at Mount Sinai, NY, NY; Icahn Genomics Institute, Black Family Stem Cell Institute, Icahn School of Medicine at Mount Sinai, NY, NY; Feil Family Brain & Mind Research Institute, Weill Cornell Medicine, NY, NY; Department of Rehabilitation Medicine, Icahn School of Medicine at Mount Sinai, NY, NY

**Keywords:** Axons, Regeneration, Circular RNA (circRNA), Spinal cord injury, CNS, RNA-seq

## Abstract

**Introduction:** Neurons transport mRNA and translational machinery to axons for local translation. After spinal cord injury (SCI), *de novo* translation is assumed to enable neurorepair. Knowledge of the identity of axonal mRNAs that participate in neurorepair after SCI is limited. We sought to identify and understand how axonal RNAs play a role in axonal regeneration.

**Methods:** We obtained preparations enriched in axonal mRNAs from control and SCI rats by digesting spinal cord tissue with cold-active protease (CAP). The digested samples were then centrifuged to obtain a supernatant that were then sequenced. We used bioinformatics analyses to identify DEGS and map them to various biological processes. We validated the DEGs by RT-qPCR and RNA-scope.

**Results:** The supernatant fraction was highly enriched for axonal mRNA. Using Gene Ontology, the second most significant pathway for all differentially expressed genes (DEGs) was axonogenesis. Among the DEGs was Rims2, which is predominately a circular RNA (circRNA) in the CNS. We show that Rims2 RNA within spinal cord axons is circular. We found an additional 200 putative circRNAs in the axonal-enriched fraction. Knockdown in primary rat cortical neurons of the RNA editing enzyme ADAR1, which inhibits formation of circRNAs, significantly increased axonal outgrowth. Focusing on Rims2 we used Circular RNA Interactome to predict that several of the miRNAs that bind to circRims2 also bind to the 3’UTR of GAP-43, PTEN or CREB1, all known regulators of axonal outgrowth. Axonally-translated GAP-43 supports axonal elongation and we detect GAP-43 mRNA in the rat axons by RNAscope.

**Discussion:** By using our method for enrichment of axonal RNA, we detect SCI induced DEGs, including circRNA such as Rims2. Ablation of ADAR1, the enzyme that regulates circRNA formation, promotes axonal outgrowth of cortical neurons. We developed a pathway model using Circular RNA Interactome that indicates that Rims2 through miRNAs can regulate the axonal translation GAP-43 a known regulator of axonal regeneration indicating that axonal mRNA contribute to regeneration.

## Introduction

Axons do not spontaneously regenerate in the injured CNS. This is in part due to diminished intrinsic capability of adult CNS neurons to regenerate axons and to extrinsic inhibition at the injury site (1,2). Neurons innervating spinal cord motor neurons, with cell bodies in the brain and brainstem and axonal projections that synapse up to a meter from the cell body, have to transport multiple cargos including mRNAs from the cell body to needed areas along the axonal shaft and at the synapse (3). Upon injury to the CNS, the neuronal cell bodies need to transport RNAs a considerable distance away to the injury site (4–9). Ribosomes are detected all along the axon and growth cone, and local translation within the axons is known to occur (9–13). The functional consequences of axonal translation are not fully known in part due to limited technologies for specifically identifying axonal mRNAs and the mechanisms by which they regulate axon growth and regeneration. Consequently, despite many studies reporting gene expression changes after spinal cord injury (SCI), there is a gap in knowledge regarding which of these expression changes occurs in descending and ascending axons responsible for voluntary movement.

The goal of this study was to define axonal mRNA profiles at the site of injury after SCI and identify the pathways modulated by differentially regulated axonal mRNAs to understand how SCI induced differentially expressed genes (DEGs) regulate axonal regeneration.

In addition to linear mRNAs, neurons express circular RNAs (14–15). CircRNAs are best characterized as post-transcriptional regulators that are well conserved in mammals, particularly enriched in synapses, and expressed during differentiation (14–17). Compared to their linear form, circRNAs are more stable (16). In the mammalian brain, circRNAs are abundant and are dynamically expressed, independent of their linear mRNA analogs (14). We are only beginning to understand circRNA function in maintaining brain physiology (14,17). For instance, the linear mRNA for *Rims2* encodes a presynaptic protein that is important for neurotransmitter release, but the function of circRims2 is still unknown (18,19). Very little is known about circRNA function and regulatory capability in neurons as a whole and if circRNA may play in axonal elongation. Elucidating the potential regulatory role of circRNAs in local axonal translation may provide a better understanding of multifacteed nature of axonal regeneration, and help in finding therapeutic targets to treat SCI.

## Methods

### SCI surgery and Locomotor Activity measurements

All animal experiments were performed according to ethical regulations and protocols approved by the Institutional Animal Care and Use Committee (IACUC) at Icahn School of Medicine at Mount Sinai and at the James J. Peters Department of Veterans Affairs Medical Center. Both institutes are Association for Assessment and Accreditation of Laboratory Animal Care (AAALAC) accredited.

Female Sprague Dawley rats 3 months of age were utilized. A moderate severity SCI at thoracic level 9 (T9) was performed using an Infinite Horizons (IH) impactor (Precision Systems and Instrumentation) (20,21). Animals were deeply anesthetized with isoflurane. A laminectomy was performed at T9. The exposed vertebral column was stabilized with clamping forceps placed rostral from T8 and caudal from T10 vertebral bodies, ensuring the exposed spinal cord is in a level horizontal plane. Control animals had laminectomy only but no contusions. The impactor tip is lowered to 3-4mms above the laminectomy site and a computer operated program is used to deliver a 250 kdyne force to the exposed spinal cord. BBB locomotor testing was done the following morning to confirm contusions was performed accurately and any animal with initial BBB Locomotor Rating score above 4 will be discarded from the study (22). Locomotor function was evaluated in a 3-foot-wide plastic tub by two independent, blinded observers from direct observation. Animals will be recorded using a GoPro camera to provide documentation of observed behaviors.

### Tissue excision for RNA-seq

24 hrs after performing the SCI, we collected tissue just rostral to the injury site which contains axotomized fibers for mRNA expression profiles. Animals were deeply anesthetized and a laminectomy at T8 was performed. With sterile scissor and forceps, about 2mm of tissue was excised just above the injury at T8. Tissue was placed into 1ml of ice-cold Miltenyi MACS Tissue Storage Solution (Miltenyi Biotech 130-100-008). Cold active protease from *Bacillus licheniformis* (Creative Enzymes NATE0633) was added (10mg/ml of this protease with 125 U/ml DNase in ice cold Miltenyi MACS Tissue Storage Solution) (23). Cells were incubated for 7 mins in a water bath set to 6°C in the cold room (23). The tissue was triturated every 2mins for 15secs. For gentle dissociation of the tissue, a Miltenyi gentleMACS at the brain program setting was ran twice in the cold room. The sample was then incubated at 6°C for an additional 8mins with trituration. Cells are then pelleted by centrifugation at 1200g for 5 mins, and supernatant was carefully collected so as not to dislodge cells. The supernatant was prepared for Bulk mRNA Sequencing. A small extract of the supernatant and cell pellet fraction were plated on glass cover slips to confirm that the supernatant was cell free and that the pellet had intact cell bodies.

### Preparation for bulk sequencing of the acellular fraction

-All supernatant fraction from the spinal cord were checked for RNA integrity by Agilent 2100 Bioanalyzer and all samples had RIN value ≥9 (24). RNA libraries for RNA sequencing were constructed using the Truseq stranded total RNA kit (Illumina Inc.) which converts the RNA in a total RNA sample into a library of template molecules of known strand origin; and purified with a RiboZero kit (Illumina). The resulting mRNA fragment inserts were then reverse transcribed to first strand cDNA using reverse transcriptase and random primer in the presence of actinomycin D. Strand specificity was achieved during second strand cDNA synthesis by replacing dTTP with dUTP in the Second Strand marking mix which includes the DNA Polymerase I and RNase H. 3’ ends were then adenylated to prevent them from ligating to each other during the adapter ligation reaction. Unique dual index adapters (i5 and i7) were then ligated allowing for greater sample indexing diversity and enables ds cDNA for hybridization onto a flow cell. The indexed double stranded cDNA was then enriched by PCR and purified to create the final cDNA library which was quantified and then loaded on the flow cell for sequencing. 50×10E6 reads per sample were obtained for Deep analysis.

### Bulk Sequencing Differential Expression Analysis Pipeline

To reduce artifacts caused by read imbalances during upper quartile normalization, we downsampled the sequencing reads of each sample to the number of reads that were detected in the sample with the lowest read counts, as described previously (24). Downsampled reads were aligned to the rat reference genome ’rn6’ using the ensemble annotation and STAR 2.5.4b (25) with the parameters set between outFilterScoreMinOverLread and outFilterMatchNminOverLread set to 0.33. Differentially expressed genes were identified with cufflinks 1.3.0 (26) (FDR 5%, minimum log_2_(fold change) = +/-log_2_(1.3)). Up-and downregulated genes were subjected separately and combined to pathway enrichment analysis using Wikipathways 2016 (27) and Gene Ontology biological processes 2018, downloaded from the Enrichr website (28), as described previously (29).

### Perfusions and sectioning of spinal cord

Fresh ice-cold 4% PFA in PBS was prepared. Rats were deeply anesthetized with Ketamine and Xylazine. We transcardially perfused with ice cold PBS and then with ice cold 4%PFA in PBS. We removed the spinal cord and post-fixed them in 30% sucrose in PBS, until they sank to the bottom of the tube. Segments were embedded in OCT. Using a Leica cryostat at -20C, we made 10μm sections of the spinal cord to be utilized for RNAscope and immunohistological analysis.

### RNA Extraction, Ribonuclease R Digestion, Reverse Transcription and qPCR

To validate circRNA using RT-qPCR, we followed the protocol from Vromman et al., 2021 (30,31), and the primer sequences we used were obtained from supplementary data on Rybak-Wolf et al., 2015 (14). Total RNA was extracted from mouse spinal cord segments (∼2 mm) rostral to the lesion epicenter 14 days after injury using TRIzol reagent (ThermoFisher) following the manufacturer’s instructions and methods previously described (21). Total RNA concentrations were determined by absorbance at 260 nm using a Nanodrop spectrophotometer (Thermo Scientific). For circRNA detection, 1000 ng of total RNA was incubated with ribonuclease R (RNAse R; Biosearch Technologies) following the manufacturer’s instructions for 20 mins at 37 C followed by a cycle of 20 mins at 65 C (30). Digested RNA was later cleaned with a RNAeasy MinElute Cleanup kit (Qiagen) and reverse-transcribed into cDNA using Omniscript reverse transcriptase (Qiagen). PowerUp SYBR Green Master Mix (Thermofisher) was used to measure mRNAs of interest by qPCR. Sequence of primers used to detect both, linear and circular forms of RIMS2, VAPA and RTN4 were as previously reported (14) for: linear RimS2 Fw: GCAAAACTACACGAGCAGCC and Rv: TCCCTGGACACTGATGGACT; circular RimS2 Fw: AAAGTCGCAGTGCCTCTCAA and Rv: TCCCATCCTGAGCGATACTTC; linear Rtn4 Fw: CAGTCCTGCCCTCCAAGC and Rv: TCAGATGCAGCAGGAAGAGC; circular Rnt4 Fw: AGATCCCTGACAGCTGTATTGT and Rv: GACGAAACAGTGTTACCTGGC; linear Vapa Fw: TGTTTGAAATGCCGAATGAA and Rv: AGTCGCTTGCACTCTTCCAT; circular Vapa Fw: TGTTTGAAATGCCGAATGAA and Rv: AGTCCTTGCACTCTTCCAT. Formation of single SYBR Green-labeled PCR amplicons were verified by running melting curve analysis. Threshold cycles (CTs) for each PCR reaction were identified by using the QuantStudio 12K Flex software. To construct standard curves, serial dilutions were used from 1/2 to 1/512 of a pool of cDNAs generated by mixing equal amounts of cDNA from each sample. The CTs from each sample were compared to the relative standard curve to estimate the mRNA content per sample; the values obtained were then normalized using peptidylprolyl isomerase A (Ppia) mRNA.

### RNAscope

We utilized RNAscope Multiplex Fluorescent v2 Assays (ACD biosciences). We follow the manufacturers protocol with minor modifications (32). Spinal Cord sections were post-fixed with 4%PFA, washed in PBS. Followed by ethanol series dehydration and subsequent rehydration series prior to treatment with hydrogen peroxide and washed again with water 3 times. Antigen retrieval was done by protease III (for cortical neurons in culture) or protease IV (for spinal cord sections) treatment in PBS for 10mins at RT; slides will then be washed twice with PBS. Sections were hybridized with probes for 2hrs at 40C then washed with wash buffer. Hybridized probes were detected using the RNA-scope Amplification (1–3) reagents for 30mins at 40C. Slides were washed and then incubated with Opal dyes (Akoya biosciences) for 15mins at 40C. Immunohistochemistry were by performed as follows. Sections were permeabilized with 0.25% Triton-X then blocked with 10% goat serum in TBS with 1%BSA overnight at 4C. Some sections were be co-labeled with antibody for β-III tubulin in TBS-BSA overnight, washed and incubated with secondary goat-anti mouse coupled to Alexa-405 for one hour at RT. Slides were washed and coverslips mounted with ProLong Gold antifade.

For Ribonuclease R digestion, after the Protease step and 3 washes with PBS, we permeabilized the tissue with 0.1% Tween-20 in PBS for 10minutes at RT. Then washed the samples with PBS three times.

### Rat primary cortical neuronal cultures

Rat primary cortical neuronal cultures were prepared as previously described with minor modifications (33,34) were dissected from postnatal day 1 Sprague Dawley rat brains, from both sexes. The cortices were incubated twice for 30mins with 0.5 mg/ml papain (Sigma) in plain Neurobasal (NB) media (Invitrogen) supplemented with DNase. At the end of the second digestion with papain, the cortices were pipetted through a 1ml serological pipette in a low volume (2 to 3 mls), to dissociate the tissue. The triturated tissue for every 3 cortices was made up to 6mls with plain NB and strained through a 70micron cell strainer. The cell suspensions were layered on an Optiprep density gradient (Sigma) and centrifuged at 1900 x *g* for 15mins at room temperature, with the brakes decreased to the lowest setting to not disrupt the gradients. The layer closest to the bottom between 1 to 2mls of a 15ml conical tube is the enriched fraction of neurons. The enriched neuronal layer was then pelleted at 1000 x g for 5 mins and counted.

### Microfluidic neurite outgrowth assay

Square microfluidic chambers (SND450, 450μm microgroove) were purchased from Xona microfluidics. The chambers were sterilized under UV for 15mins and soaked in 70% ethanol for 2mins and allowed to air dry under a sterile TC hood. We used MatTek dishes (P50G-1.5, MatTek corp.) that we pre-coated with PLL overnight and then rinse 3 times with sterile water and air dried overnight in a TC BSL2 hood. Using autoclaved forceps we carefully place the microfluidic chamber on the glass area of the MatTek dish, and gently apply pressure to ensure the chambers are sitting on the glass. Primary cortical neurons were diluted to 5×10E6 cells/ml in NB supplemented with B27, L-glutamine and antibiotics, and in a final volume of 200µls. Approximately 15μls of the cell suspension was carefully inserted into the top well of one side of microfluidic chamber. We then placed the Mat-tek dishes with the microfluidic chambers inside a 37°C incubator for 20mins to allow the neurons to adhere. All wells were filled with 150µls of supplemented NB. Once we observe neurites growing across the microgroove (3-4days), we then performed an axotomy.

To quantify the outgrowth we immunostained using a monoclonal antiβIII tubulin antibody (Tuj1;Covance) and Alexa Fluor -405-(Blue) or 488-(Green) conjugated anti-mouse IgG (Invitrogen). For quantification, images were taken, and the length of the longest neurite for each neuron was measured using MetaMorph software (Molecular Devices). All imaging was performed on a Zeiss LSM 880 confocal microscope.

### siRNA transfection

All siRNA were purchased from Accell smartpool siRNA (Thermo Scientific), following the manufacturers protocol. We targeted ADAR1 (Rat Accession number XM_006232778.3), scrambled non-targeting siRNA was also utilized for controls. Cortical neurons were plated on one side of the microfluidic chambers. We waited 2-4 hrs after plating to allow the cells to adhere. Utilizing 1µM siRNA in 150µls of Accell Delivery Media was added to the neuronal cell bodies and incubated overnight at 37C incubator. The next morning, we added 150µls of supplemented NB. We incubated for a further 72hrs before stopping the experiment with 4% paraformaldehyde and 4% sucrose. We then stained the chambers with β-III tubulin (Alexa-488) and in individual neurons panel, we also used Actin-Red staining (Thermo Scientific) and quantified the neurite length.

### Statistical analyses

All analyses were performed using GraphPad Prism software, and data are represented as mean ± SEM. Statistical significance was assessed using paired one-tailed Student’s *t* tests to compare two groups, and one-way ANOVAs with Bonferroni’s *post hoc* tests to compare between three or more groups.

## Results

We adapted a tissue dissociation protocol used for single cell RNAseq (scRNAseq) that uses cold active proteases to gently dissociate tissue and isolate cells (23). Low speed centrifugation supernatant of the dissociated cell suspension was used to pellet cells and recover the supernatant. This was followed by bulk RNA sequencing (RNAseq) of the supernatant to identify mRNAs that are more associated with the acellular fraction of cells. We identified the DEGS by comparison of RNAseq data for control vs SCI groups (21,35–36). The DEGS are shown in Table 1A. We detected 3 known axonal microtubule associated protein (MAP) mRNAs, MAP4, Tau (MAPT) and MAP7 and all were significantly down-regulated after SCI. MAP4 was the most abundant axonal mRNA detected. Several axonal RNA transport mRNAs were detected. RBP1 is up-regulated nearly 15-fold after SCI (37,38). We detected RNAs that function in the molecular motors that drive anterograde transport (Kinesins such as KIF5A), and retrograde transport (Dynein, DYNC1LI2) (39). Other mRNAs detected included those encoding Guanine Deaminase (GDA), MACROH2A1, IMPACT and Rims2.

**Table 1A:** DEG analysis indicates enrichment for axonal RNA. Axonal markers DEGS detected when comparing uninjured rat spinal cord (No-SCI) to ones that received contusions to the spinal cord (SCI)., P-Adj (Q-) values adjusted for multiple comparison testing and Fold changes are shown. Significant Q-values are in Bold and Italic type. Fold changes are presented as Log2 fold changes. We made a short list of both novel and known DEGs (green font). For our Bulk RNA-seq analysis we read 50×10E6 reads per sample, giving us the ability to detect low abundance transcripts. Bold and italic numbers for P-Adj indicate significant change.

| **DEG** | **No-SCI** | **SCI** | **P-Adj** | **Log2 Fold** |
| --- | --- | --- | --- | --- |
| <u>**Axonal Markers**</u> |  |  |  |  |
| <u>*Axonal MAPs*</u> |  |  |  |  |
| MAP4 | 20198.17 | 6958.2 | ***0.014066*** | -1.537438 |
| MAPT (tau) | 2223 | 1231.45 | ***0.013229*** | -0.8521476 |
| MAP7 | 712.95 | 220.02 | ***3.92E-13*** | -1.695605 |
| <u>*Axonal mRNA Transport*</u> |  |  |  |  |
| RBP1 | 92.79 | 1467.37 | ***1.81E-12*** | 3.983084 |
| <u>*Axonal Anterograde*</u> |  |  |  |  |
| KIF5A | 36.6375 | 15.52 | ***1.79E-02*** | -1.239592 |
| <u>*Axonal Retrograde*</u> |  |  |  |  |
| DYNC1LI2 | 1595.324 | 570.756 | ***6.66E-08*** | -1.482903 |
| <u>**DEGs**</u> |  |  |  |  |
| GDA | 6.74 | 51.19 | ***1.07E-06*** | 2.926016 |
| NGF | 2.77 | 12.18 | ***0.001586*** | 2.137659 |
| MACROH2A1 | 243.62 | 607.32 | ***0.026033*** | 1.31786 |
| NCAM1 | 202.47 | 82.65 | ***2.51E-06*** | -1.292713 |
| IMPACT | 22.42 | 8.6 | ***0.0298*** | -1.382178 |
| Rims2 | 183.564 | 68.53 | ***0.00026*** | -1.421475 |

**Table 1B:** DEG analysis suggests minimal glial cell contamination. The table lists DEGs of known dendritic and glial cell markers. Adjusted P values, corrected for multiple comparison testing and fold changes are shown. Bold numbers indicate significant change.

| **DEG** | **No-SCI** | **SCI** | **P-Adj** | **Log2 Fold** |
| --- | --- | --- | --- | --- |
| <u>Dendritic Markers</u> |  |  |  |  |
| MAP2 | 2139.92 | 859.51 | **0.000225** | -1.315974 |
| <u>Astrocytic Markers</u> |  |  |  |  |
| GFAP | **Not Detected** |  |  |  |
| S100B | **Not Detected** |  |  |  |
| EAAT1/GLT1 | **Not Detected** |  |  |  |
| EAAT2 | **Not Detected** |  |  |  |
| AQP4 | **Not Detected** |  |  |  |
| ALDH1L1 | 8.19166 | 3.01872 | **3.32E-05** | -1.44022 |
| ALDOC | **Not Detected** |  |  |  |
| SLC1A3 | **Not Detected** |  |  |  |
| <u>Microglia Markers</u> |  |  |  |  |
| Iba1 | **Not Detected** |  |  |  |
| P2YR12 | **Not Detected** |  |  |  |
| TMEM119 | 8.66 | 6.49 | 0.659135 | -0.4151374 |
| SPP1 | 221.958 | 265.43 | 0.703813 | 0.2580448 |
| TREM2 | 21.771 | 32.834 | 0.516 | 0.5927786 |
| AXL | 39.485 | 22.511 | 0.276637 | -0.8107108 |
| LPL | 3.096 | 1.752 | 0.07306 | -0.8213534 |
| CST7 | 3.682 | 6.166 | 0.478 | 0.7437049 |
| <u>Oligodendrocyte</u> | <u>Markers</u> | <u>Associated</u> | <u>with cell bodies</u> |  |
| MBP | **Not Detected** |  |  |  |
| APC | 185.118 | 111.813 | 0.052068 | -0.727355 |
| NG2 | **Not Detected** |  |  |  |
| O4 | **Not Detected** |  |  |  |
| MOG | 138.127 | 63.8503 | 0.137524 | -1.11323 |
| <u>Oligodendrocyte</u> | <u>Markers</u> | <u>Associated</u> | <u>with cell</u> | <u>processes</u> |
| CNP | 1887.484 | 614.932 | **0.016289** | -1.61797 |
| RIP | **Not Detected** |  |  |  |
| GALC | 10.114 | 6.4 | 0.528041 | -0.66017 |
| OMGP | **Not Detected** |  |  |  |
| MAG | 271.936 | 87.3568 | **9.69E-06** | -1.63828 |

**Table 1C:** DEG analysis suggests minimal spinal interneuron RNA. DEGs of spinal interneuron markers. Adjusted P values and Fold change are shown. Additional markers can be found in Supplementary Table 1.

| **DEG** | **No-SCI** | **SCI** | **P-Adj** | **Log2 Fold** |
| --- | --- | --- | --- | --- |
| *<u>Interneurons</u>* |  |  |  |  |
| ASCL1 | 2.01347 | 2.88741 | 0.51936 | 0.520092 |
| BARH1 | **Not Detected** |  |  |  |
| BRN3A | **Not Detected** |  |  |  |
| BRN3B | **Not Detected** |  |  |  |
| CBLN2 | 6.23645 | 2.839465 | 0.162549 | -1.135106 |
| DRG11 | **Not Detected** |  |  |  |
| GAD2 | 2.78046 | 1.833779 | 0.183802 | -0.6005038 |
| GLYT2 | **Not Detected** |  |  |  |
| GSH1 | **Not Detected** |  |  |  |
| GSH2 | **Not Detected** |  |  |  |
| ISL1 | **Not Detected** |  |  |  |
| LHX1 | **Not Detected** |  |  |  |
| LHX2 | **Not Detected** |  |  |  |
| LHX5 | **Not Detected** |  |  |  |
| LHX9 | **Not Detected** |  |  |  |
| LMX1B | **Not Detected** |  |  |  |
| MATH1 | **Not Detected** |  |  |  |
| NPY | 302.955 | 221.5497 | 0.700791 | -0.4514731 |
| PAX2 | **Not Detected** |  |  |  |
| PTF1A | **Not Detected** |  |  |  |
| RORA | 52.0532 | 35.422 | 0.526817 | -0.5553411 |
| RORB | 9.1919 | 6.02137 | 0.4115 | -0.6102714 |
| SMARCA2 | 7.78941 | 7.35343 | 0.927974 | -0.08309677 |
| TAG-1 | **Not Detected** |  |  |  |
| TLX-3 | **Not Detected** |  |  |  |
| VGLUT2 | **Not Detected** |  |  |  |

**Table 1D:**
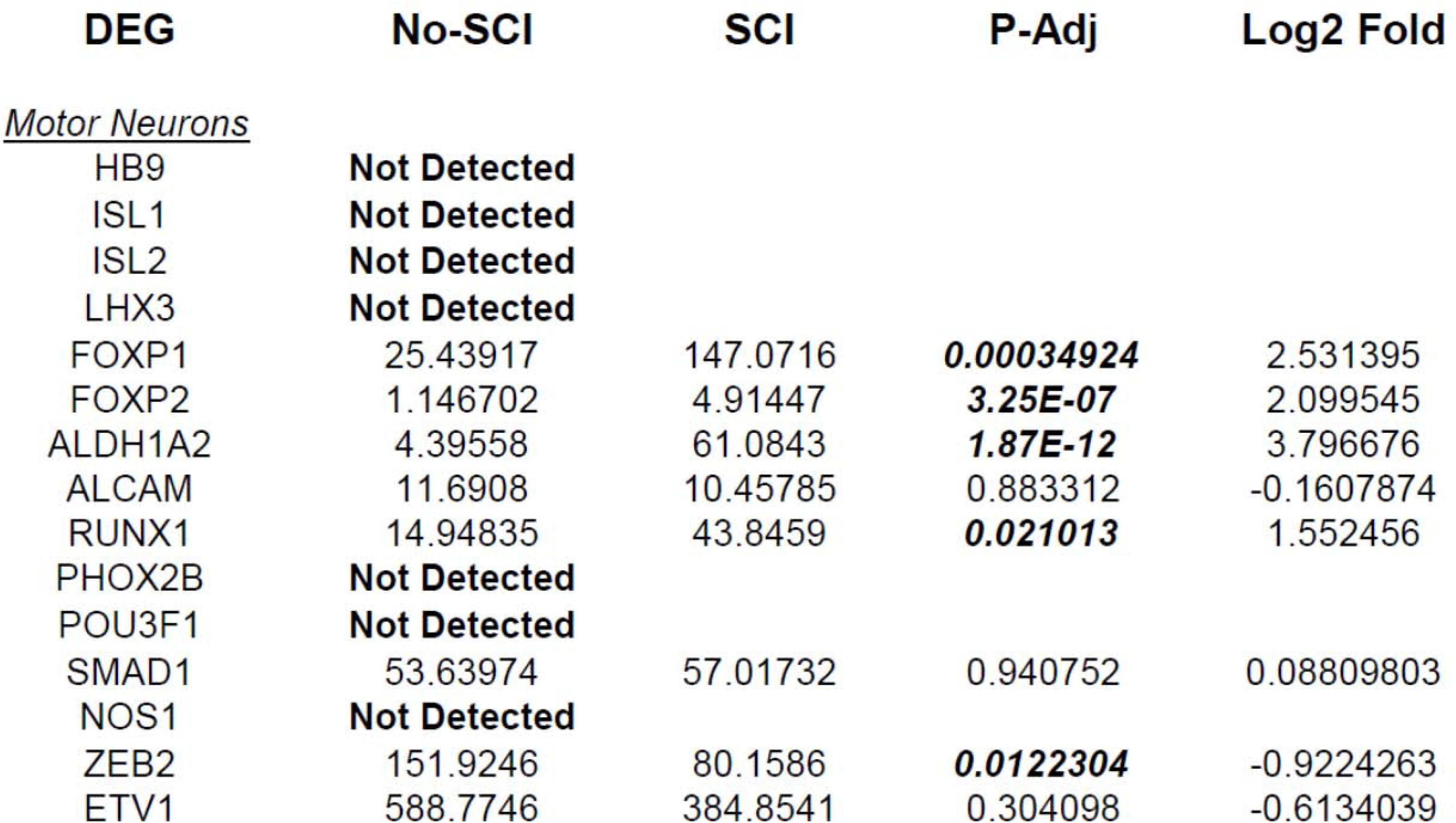
DEG analysis for Motor Neuron RNA. The table lists motor neuronal marker DEGs. Adjusted P-value and fold change are given. Bold numbers indicate significant changes Supplementary Table 1.

| DEG | No-SCI | SCI | P-Adj | Log2 Fold |
| --- | --- | --- | --- | --- |
| *Motor Neurons* |  |  |  |  |
| HB9 | Not Detected |  |  |  |
| ISL1 | Not Detected |  |  |  |
| ISL2 | Not Detected |  |  |  |
| LHX3 | Not Detected |  |  |  |
| FOXP1 | 25.43917 | 147.0716 | **0.00034924** | 2.531395 |
| FOXP2 | 1.146702 | 4.91447 | **3.25E-07** | 2.099545 |
| ALDH1A2 | 4.39558 | 61.0843 | **1.87E-12** | 3.796676 |
| ALCAM | 11.6908 | 10.45785 | 0.883312 | -0.1607874 |
| RUNX1 | 14.94835 | 43.8459 | **0.021013** | 1.552456 |
| PHOX2B | Not Detected |  |  |  |
| POU3F1 | Not Detected |  |  |  |
| SMAD1 | 53.63974 | 57.01732 | 0.940752 | 0.08809803 |
| NOS1 | Not Detected |  |  |  |
| ZEB2 | 151.9246 | 80.1586 | **0.0122304** | -0.9224263 |
| ETV1 | 588.7746 | 384.8541 | 0.304098 | -0.6134039 |

In Table 1B, we show detection of the dendritic marker MAP2, but its levels are over 10-fold lower than the axonal MAPs detected in Table 1A. We detected none of the known astrocytic markers, Glial fibrillary acidic protein (GFAP), S100B, Excitatory amino acid transporter 1 and 2 (EAAT1 and 2) and aquaporin 4 (AQP4), except for ALDH1L1 at low levels of expression (40–42). Prominent microglial markers, such as Iba1 and P2YR12, were not detected, suggesting that the supernatant enriched in axonal mRNAs was relatively free from other cell contaminants (42–43). Several microglial cell markers were detected at low levels, but there was no difference in their P-Adj value when comparing controls and SCI (42–48). In general, the number of transcripts associated with microglial markers was relatively low (<100), whereas there were >10,000 axonal-associated markers detected. We have mixed results for detection of oligodendrocyte markers, but only CNP and MAG were significantly altered by SCI (CNP) which are known markers for the oligodendrocyte processes that myelinate axons, suggesting that we are detecting processes associated with the axons and not the oligodendrocyte cell bodies (49,50). Taken together, the robust axonal signals that are associated with axonal markers with minimal dendritic and glial cell contamination, indicate that we had enriched the supernatant fraction for axonal RNA (Figure 1).

**Figure 1.**
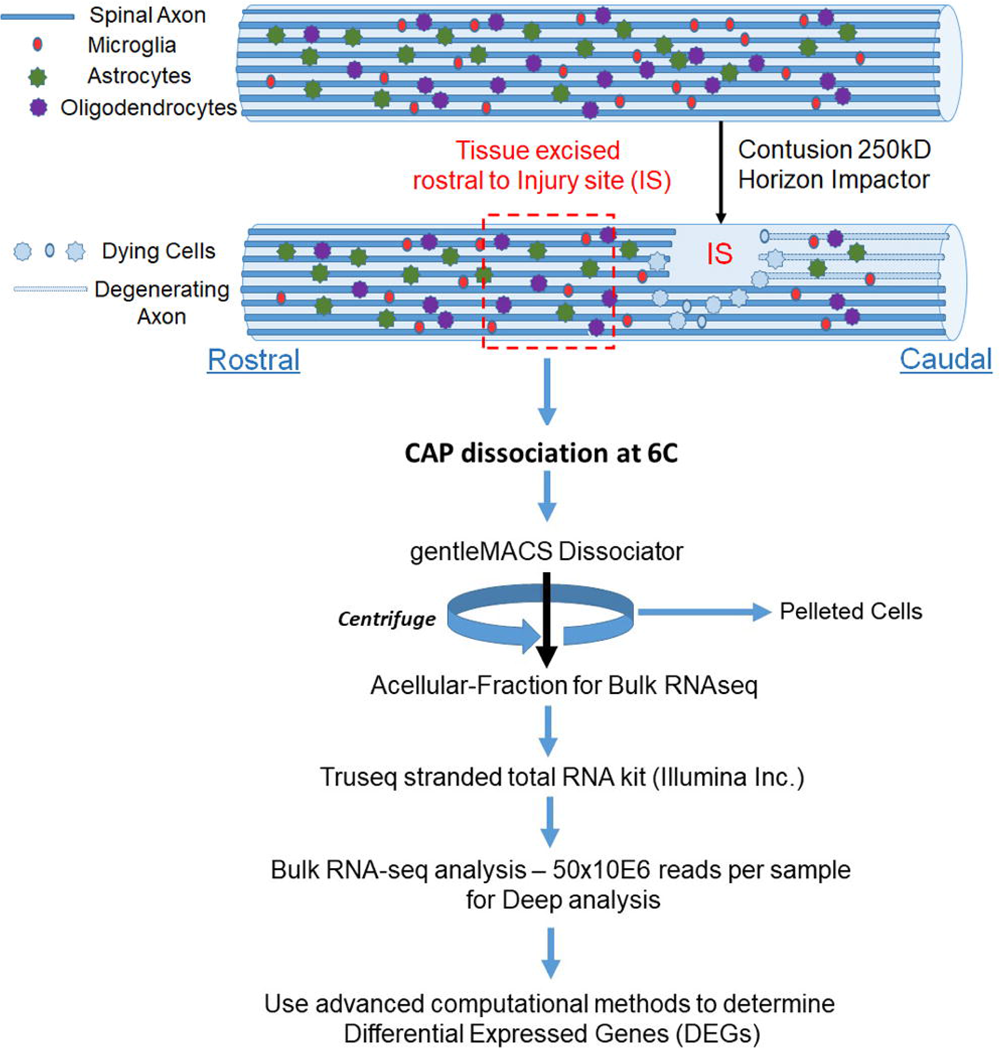
Schematic for enrichment of acellular RNA from the spinal cord. Adult Sprague Dawley rats have laminectomies at T9 and performed contusions using an Infinite Horizons impactor with a force of 250kdyne to induce SCI, while control animals have laminectomy only. We wait 24hrs and then excised tissue approximately one lamina rostral from the injury site (IS) and treated with CAP at 6^0^C. We used the gentleMACS Dissociator in the cold room set to the Brain setting and then centrifuged the sample at 1200RPM for 5mins. We used the supernatant fraction for Bulk RNA-seq. We identified DEGs between laminectomy alone vs. SCI. We used Gene ontology to identify biological processes (pathways) from DEGs

**Figure 2.**
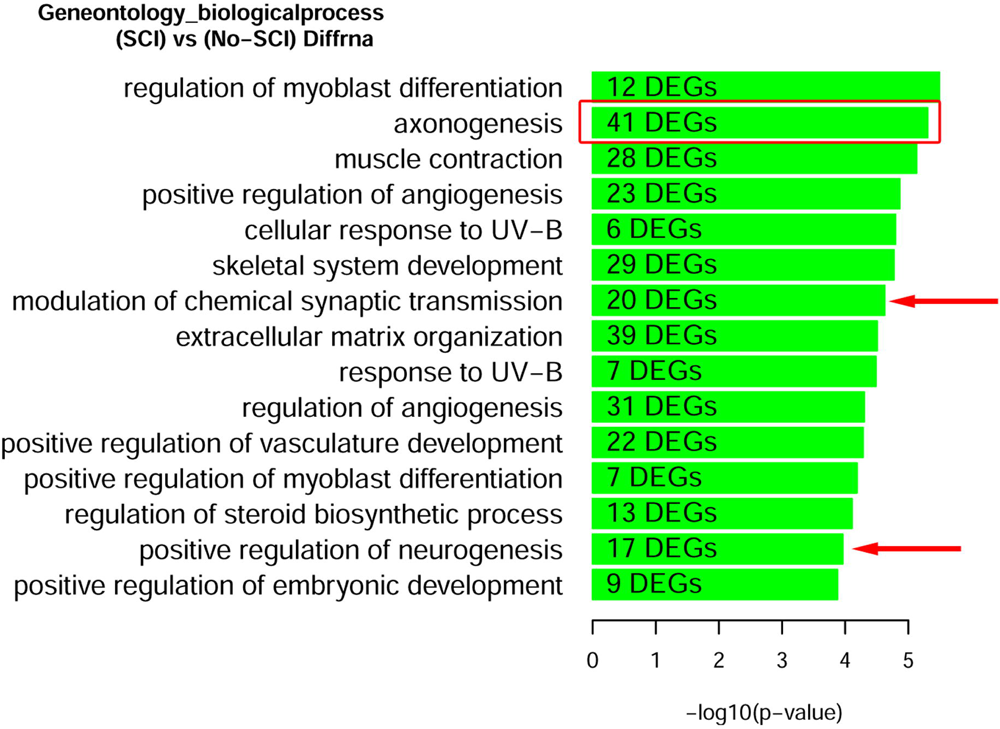
Gene Ontology enrichment of DEGs after SCI identifies that axonogenesis is a robustly significant biological processes. Rats with and without spinal cord injury (SCI) were subjected to CAP purification for the acellular fraction for Bulk RNA-seq analysis. Lists of DEGs were used for enrichment analysis to predict differentially regulated pathways,. We find the second most significant pathway to be for Axonogenesis (red box). Other pathways relevant to the CNS are demarcated with a red arrow. A complete list of Up-regulated and Down-regulated GO-pathways is shown in Supplementary Figure 1.

We also looked for known markers of Spinal Interneurons (Table 1C and in Supplementary Table 1). Only a few (ASCL1, CBLN2, GAD2, RORA, RORB and SMARCA2) of the interneuron sub-type specific marker RNAs were detected. These had low abundance and were not significantly altered by SCI. We also looked for motor neuron markers and found that roughly half of them were undetected. For the remaining mRNA species for motor neurons, 5 were significantly altered by SCI and 4 of the motor neuronal markers (RUNX1, ZEB2, FOXP1 and FOXP2) have also been reported to be either in the axons or involved in mediating axonal elongation (51–54). Overall, our analysis indicates that we do have enrichment of axonal RNA (see Table1A-D & Supplementary Table1) with minimal contaminating glial cells, spinal interneurons, or motor neurons.

We validated the expression of selected DEGs in axons of cultured primary rat cortical neurons plated in microfluidic chambers using RNA-scope. RNAscope enables us to visualize single RNA molecules within a cell using confocal laser microscopy. One novel DEG detected was GDA which can promote microtubule assembly in neurons, a necessary step for axonal extension, which is up-regulated after SCI (55). In Figure 3, we show validation of the RNAseq detected DEGs using RNAscope. We also probed for GAP-43, which was known to be located in the axons (7). We observed that GAP-43 (green) and GDA (red) were present in the cell bodies (Figure 3A left side and 3H), but 24 hrs post axotomy, the expression of both GAP-43 and GDA (Figure 3H) was increased compared to no axotomy control. Elevated GAP-43 expression was anticipated, as it is known to be elevated in neurons after CNS injury (56,57). In addition to the increased axonal mRNA levels, both GAP-43 and GDA mRNAs are transported to more distal axonal regions following axotomy (see magnified region in Figure 3D, quantification in Figure 3F&G). The graphs in Figure 3G&H are the average of 4 independent experiments for which RNA was quantified blindly by two independent reviewers. We also examined other DEGs, such as NGF and IMPACT. NGF has a well established role in growth support of neurons and promoting neurite outgrowth and axonal extension (58). IMPACT is found in the CNS, having a role as a translational regulator (59). In Figure 3E we show an axon exiting the microgroove and both NGF (red) and IMPACT (green) were detected but they were localized to different regions of the axons. We further validated additional DEGs detected in both cortical neurons and in adult rat spinal cord. We detected MACROH2A1 (red) and MAP4 (green) in cortical neurons in microfluidic chambers and in sectioned uninjured rat spinal cord by RNAscope (Figure 4A and B) and found their movement increases after axotomy. MACROH2A1 is a variant of histone H2A; its depletion in mouse brains has been shown to boost hippocampal synaptic plasticity (60). The RNA for MACROH2A1 (red) and MAP4 (green) by RNAscope (Figure 4B), localized in the white matter outside of the DAPI (blue) stained nuclei. In a sagittal section of the rat spinal cord, we immunostained for β-III tubulin (blue) and detected the mRNA for MACROH2A1 (red) by RNAscope (Figure 4C), that co-localized with the β-III tubulin. Since we are interested in axonal translation, we also confirmed the expression of S6 Ribosomal Protein in the growing axons by immunohistochemistry (Supplementary Figure 3). Ribosomes were detected in the growth cone tip detected by Actin (red, see arrow) and along the axonal shaft with β-III Tubulin.

**Figure 3.**
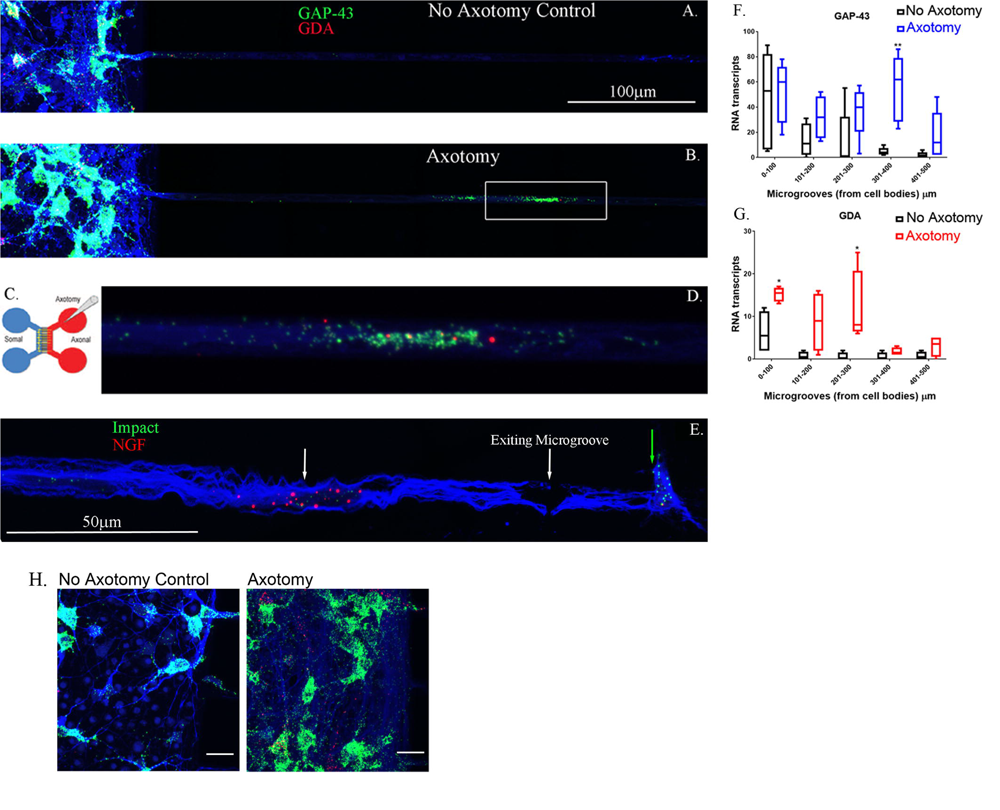
DEGs found from the RNA-seq data detected in growing axons using RNAscope. Primary rat cortical neurons were plated in microfluidic chambers and given 4 days to put axons across the microgrooves and into the axonal chamber. On day 4 we performed an axotomy and waited 24 hrs (see Cartoon in **C**.), where we sever the growing axons from the somal compartment. Comparing no axotomy controls (**A**.) to lesioned axons **(B**.) we see that in the cell body side (somal) on the left, there is in increase in GAP-43 (green) in the neuronal cell bodies that had axotomy in B. We also observed in the microgrooves of the lesioned axons, there was more abundant GAP-43 and GDA (red) in the microgrooves. The white box in B. is shown in higher magnification in **D.**, displaying the GAP-43 and GDA RNA. The axons are observed by immunohistochemical staining for β-III tubulin which is shown in blue. **E**. Using RNAscope for Impact (Green) and NGF (Red), and looking in the microgrooves as the axons are exiting from them, we see a spatial distribution of NGF in the axons (see white arrow) comparing to Impact (see green arrow) which is found at the growing tip of the axons**. F**. Quantification of GAP-43 RNA in 100 micron increments within the microgroove with and without axotomy. The graph is the average of 4 independent experiments. **G.** Quantification of GDA RNA in 100 micron increments within the microgroove with and without axotomy. The statistics is t-test with *p<0.05 and **p<0.01. H. Closeup image of the cell bodies with and without axotomy and probed by RNAscope for GAP-43 (green) and GDA (red).are shown As predicted from MRNA-Seq data both GAP-43 and GDA should be elevated after axotomy and this is what we observe. The neurons are also immunostained with β-III tubulin which is shown in blue.

**Figure 4.**
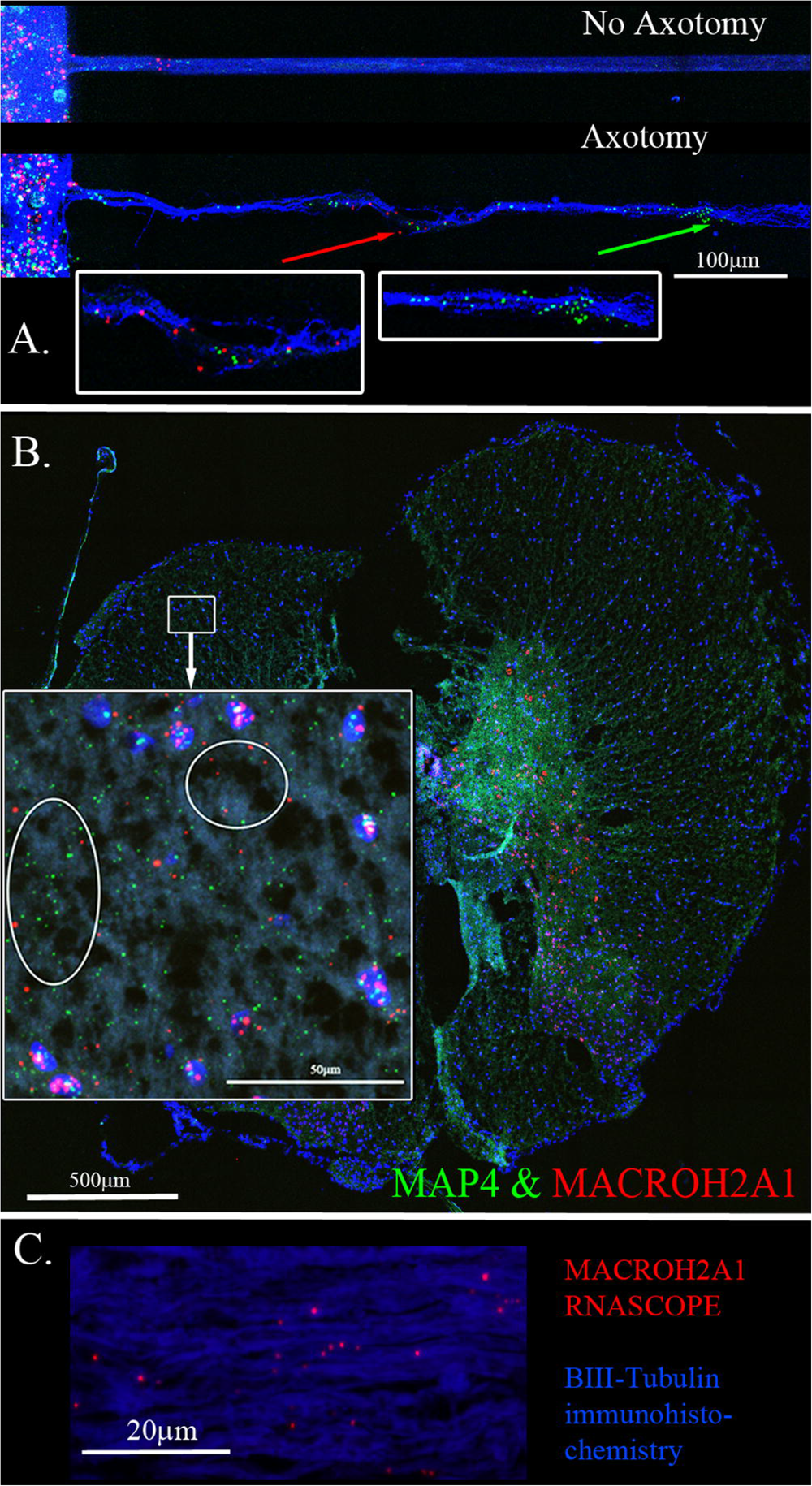
MACROH2A1 and MAP4 RNA are detected in axons with axotomy and in the white matter of rat spinal cord. **A.** Cortical neurons in microfluidic chambers without axotomy have little detectable MACROH2A1 (Red) and MAP4 (Green) by RNAscope in growing axons immunostained with β-III tubulin (Blue). After Axotomy both multiple transcripts of MACROH2A1 (red arrow) and MAP4 (green arrow), several hundred microns further away when comparing to No Axotomy. **B**. Both MACROH2A1 and MAP4 transcripts are detected by RNAscope in the white matter of the uninjured rat spinal cord in the transverse plane, cell nuclei are stained with DAPI (blue). Looking at the magnified region in the white box, both MACROH2A1 and MAP4 transcripts are detected outside of DAPI positive nuclei and in the white matter. **C.** Sagittal section of spinal cord revealing axons are immunostained with β-III tubulin (Blue) and RNAscope detection of MACROH2A1 (Red), revealing the presence of this RNA transcript along the axons.

Another DEG decreased after SCI was Rims2 which was downregulated significantly. Rims2 is reported to be predominately a circular RNA (circRNA) in the rodent brain (14). We found over 200 putative circRNAs in our RNAseq data. In Table 2 we list the top 20 most significantly altered putative circRNAs found. Those above the black line were down-regulated while those below the line were up-regulated. We confirmed by RT-qPCR in combination with digestion by ribonuclease R (RbnR, an enzyme that can only digest linear mRNAs but not digest circRNAs), that we could detect the circRNAs for Rims2, RTN4 and VAPA in mouse spinal cord tissue with and without contusion (Figure 5). Supporting the RNA-seq analysis, RT-qPCR for both Rims2 and RTN4 confirmed that their mRNAs were significantly downregulated after SCI, while VAPA was up-regulated.

**Figure 5.**
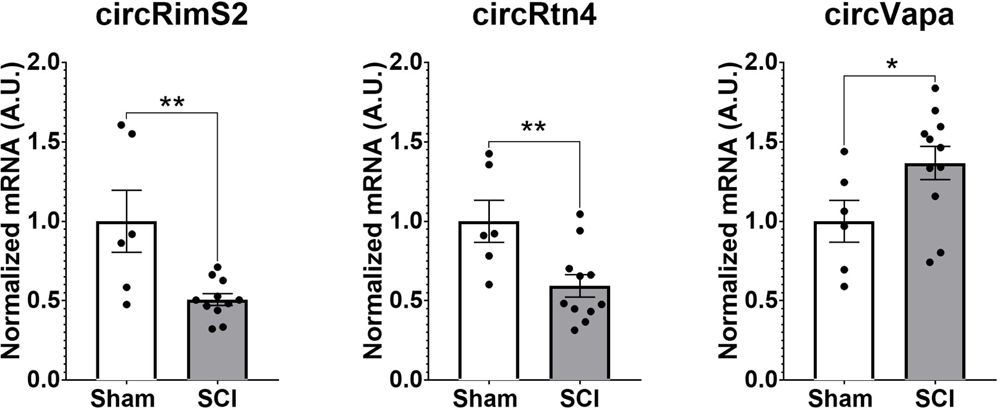
Validation by RT-qPCR in combination with RbnR treatment for identification of circRNA from mouse spinal cord. Using mouse spinal cord from animals with (N=11) and without contusions (N=6) at 14 days post injury, we combined RbnR treatment and RT-qPCR for detection of circRims2, circRtn4 and circVAPA. We found that the trends were the same as in the RNA-seq dataset, in that both circRims2 and circRtn4 were down-regulated while circVAPA was up-regulated. CircRNA changes are significant as determined by t-test with **p<0.01 and *p<0.05.

**Table 2.** List of putative circRNA detected by our RNA-seq. From our analysis, we have detected over 200 putative circRNA. We made a short list of the 20 with the most significant P-Adj values. The circRNA above the line are down-regulated after SCI and the ones below are up-regulated.

| **DEG** | **No-SCI** | **SCI** | **P-Adj** | **Log2 Fold** |
| --- | --- | --- | --- | --- |
| <u>circRNA</u> |  |  |  |  |
| GRM5 | 23.29577 | 2.694973 | **7.65E-07** | -3.111725 |
| GRM3 | 102.725 | 18.6941 | **4.31E-08** | -2.458133 |
| SYN1 | 30.3644 | 6.49934 | **0.0001** | -2.224016 |
| MAGI1 | 214.236 | 52.34484 | **0.0006** | -2.033082 |
| ZKSCAN1 | 30.0316 | 8.962 | **0.01046** | -1.744589 |
| ASPH | 104.0453 | 36.66751 | **1.07E-05** | -1.504637 |
| RIMS2 | 183.564 | 68.53002 | **0.00026** | -1.421475 |
| TRIM2 | 324.4108 | 130.2952 | **0.00255** | -1.316038 |
| NCAM2 | 7.72297 | 3.106897 | **0.00158** | -1.313681 |
| NCAM1 | 202.4714 | 82.64531 | **2.51E-06** | -1.292713 |
| RTN4 | 137.3176 | 53.3194 | **0.00477** | -1.364784 |
| FOXO3 | 25.1336 | 59.5538 | **0.00924** | 1.244576 |
| TCF12 | 33.516 | 85.27979 | **0.00676** | 1.347501 |
| VAPA | 171.73 | 445.116 | **0.04151** | 1.374039 |
| PRRX1 | 24.0978 | 63.6758 | **0.00119** | 1.401844 |
| UCK2 | 18.7675 | 64.8097 | **1.43E-07** | 1.787973 |
| COL3A1 | 2.31731 | 11.5008 | **2.20E-06** | 2.311211 |
| FOXP1 | 25.43917 | 147.0716 | **0.00034924** | 2.531395 |
| GDA | 6.73558 | 51.191 | **1.07E-06** | 2.926016 |
| CLMP | 1.574086 | 15.92662 | **3.56E-11** | 3.338854 |

To determine if circRNAs are involved in axonal regeneration, we knocked down the RNA editing enzyme ADAR1 by siRNA in primary rat cortical neurons (61). ADAR1 antagonistically regulates the formation of circRNAs (61). ADAR1 ablation led to increases in axonal outgrowth (Figure 6). We confirmed by RT-qPCR that ADAR1 siRNA was decreasing ADAR1 expression (over 60% reduction, see Supplementary Figure 2A). Controls (with scrambled siRNA) grew axons up to a maximum of 900μm from the edge of the microgrooves. ADAR1 knockdown resulted in longer (1.2mm from the microgroove) and more abundant outgrowth (Figure 6B) at 800 and 900μm from the edge of the microgrooves.

**Figure 6.**
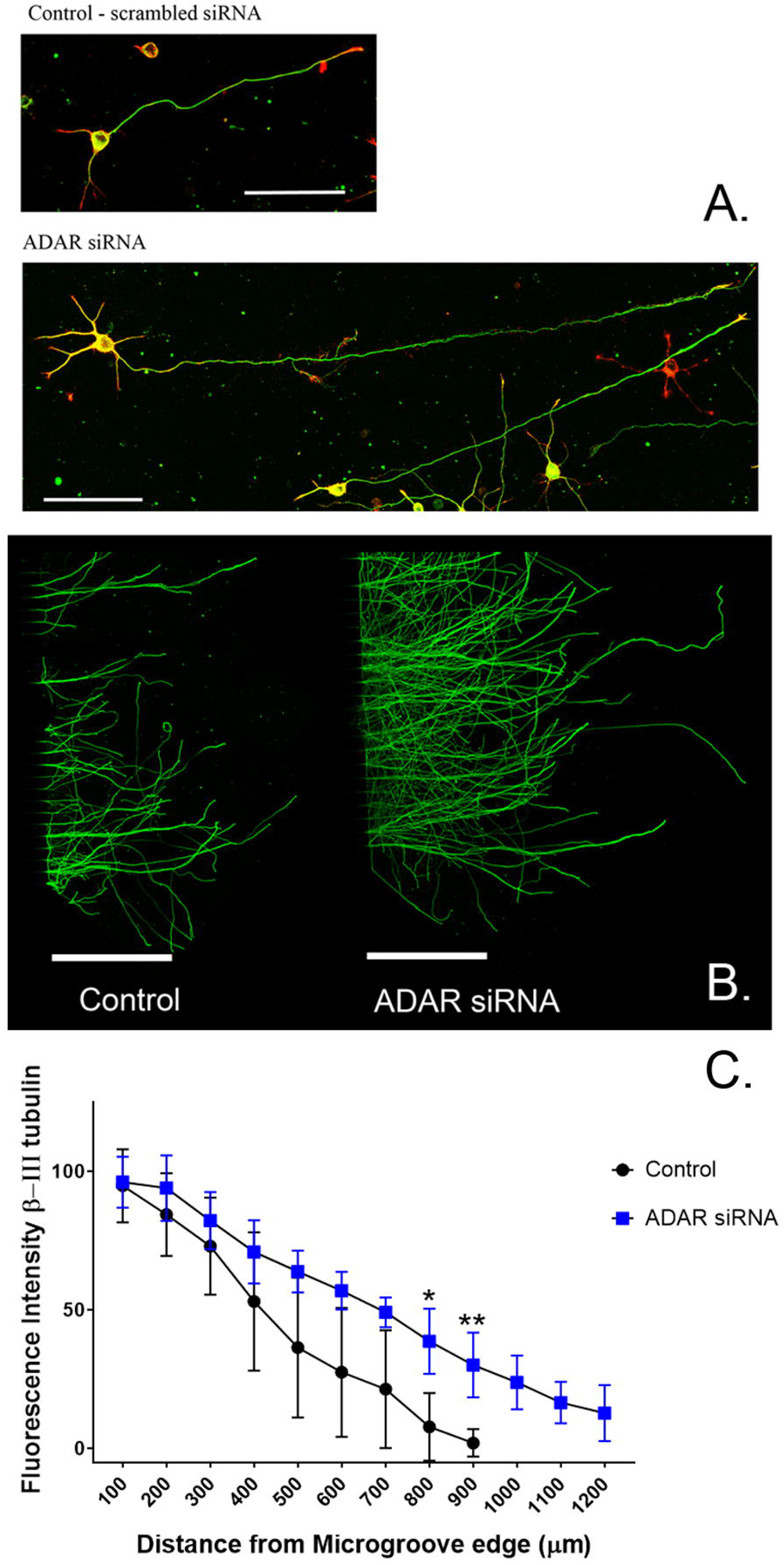
Treatment with siRNA against ADAR1 compared to scrambled controls promote longer axons in rat primary cortical neurons. A. Axons were stained with β-III tubulin (green) antibodies and actin (red) stain. B. Bottom panel, Cortical neurons in microfluidic chambers longer axons with ADAR 1 siRNA compared to scrambled controls, Scale right panels 20 μm and microfluidic chambers in left panel 500μm. C. The graph is the average of 4 independent experiments in the microfluidic chambers and data points are the average with standard deviation. Statistics is t-test (*p<0.05 and **p<0.01) comparing Control to ADAR siRNA (blue) at a fixed distance in the microgrooves. At 800 and 900 microns, ADAR siRNA (blue) were growing significantly longer compared to controls. The graph is also displayed in Supplementary Figure 2, where we show the distribution of points for each condition.

We confirmed the presence of circRims2 in rat spinal cord (Figure 7). Adjacent sections of uninjured spinal cord from the same animal were used for RNAscope detection of GAP-43 (green) and Rims2 (red) transcripts with β-III tubulin (blue) immunohistochemistry. One section was treated with ribonuclease R (right image) to eliminate linear transcripts (GAP-43) and linear Rims2; the adjacent section was used as an untreated control. Both transcripts were present in untreated sections; with ribonuclease R treatment, GAP-43 expression was absent, but Rims2 was detected suggesting that it is circular and therefore protected from the ribonuclease.

**Figure 7.**
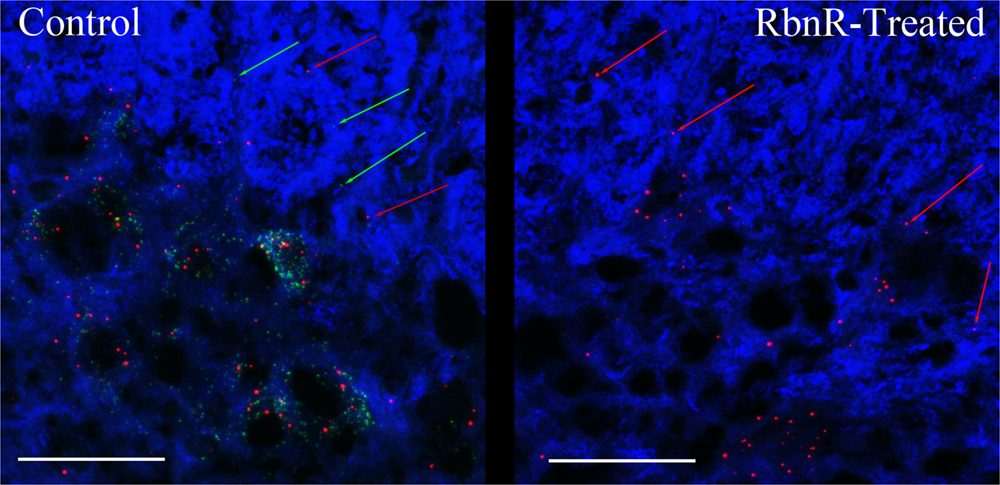
circRims2 is detected in rat spinal cord by RNAscope. Uninjured spinal cord section probed by RNAscope for GAP-43 (green) and Rims2 (red) transcripts and for with β-III tubulin (blue) immunohistochemistry. Adjacent sections were processed identically except that one was treated with RbnR to remove all linear transcripts, such as GAP-43 and preserve circRNAs. In the control (left) we detect both GAP-43 (green arrows) and Rims2 (red arrows). With RbnR-treatment, only the circRims2 remains and GAP-43 is undetected (right).

Since the known roles for circRNA are to serve as miRNA ‘sponges’ or to bind to RNA-binding proteins (RBP) (15–17), we took a computational approach by using Circular RNA Interactome to determine which miRNA or RBP could be binding to circRims2 (62). We detected several miRNAs that have been implicated in regulating axonal outgrowth, such as miR-155 (63,64), miR-21 (65) and miR-377 and -192 that both target GAP-43 (66,67). From the Circular RNA Interactome analysis, we hypothesize a RNA pathway in axons from circRims2 to GAP-43 mRNA that can regulate the local levels of GAP-43 protein (62). Additionally, we find that circRims2 can interact with the RBP, FUS, which is involved in neurite outgrowth and axonal transport (Supplementary Figure 4) (70–73).

## Discussion

Identifying the axonal RNAs transported to the site of axotomy in the injured spinal cord is critical to determining which of these may be required for promoting axonal repair. A reliable technique for determining specific axonal RNA without contaminating glial cell or spinal neuronal cell bodies RNA has been lacking. Here, we report a simple procedure for enriching axonal fractions of spinal cord incorporating cold active proteases that function at 6°C that helps reduce artifacts from transcription or translation of RNA that can occur with enzymatic digestion at 37°C (Figure 1) (23). Our analysis indicates the enrichment of axonal RNA (see Table1A-D & Supplementary Table1). Intriguingly, the only significant DEG for oligodendrocytes detected were for CNP and MAG, which are known to be associated with myelinating processes that enwrap axons (49,50). Though no significant spinal interneurons markers were detected, we detected five motor neuron (MN) marker RNAs that were significantly modulated when comparing control preparations to SCI. In the literature, 4 of these 5 mRNAs (RUNX1, ZEB2, FOXP1 and FOXP2) have also either been detected in the axons or are involved in axonal outgrowth (49–52) and hence we cannot rule out that they are axonal in origin. We validated that the DEGs detected by RNAseq could be detected in the growing axons of cortical neurons in culture and in the white matter of the adult rat spinal cord. Similar to what has been reported for GAP-43 elevation after axon injury, we observed elevated GAP-43 levels after axotomy of cortical neurons (Figure 3H) (56,57). In our RNA-seq dataset, we detected a novel DEG, GDA, which is found at synapses, and which increases significantly after injury (55). We confirmed by RNA-scope that not only do GDA levels go up after axotomy but GDA transcripts travel significantly further down the axons growing in the microgrooves (Figure 3). Similarly, MACROH2A1, which is elevated after SCI, and we detect it in the axons of cortical neurons as well as in the white matter of the spinal cord outside of the neuronal cell bodies (Figure 4). These findings support the conclusion that we are indeed detecting axonal localized RNAs.

One DEG in the axon, Rims2, has been found to be predominately a circRNA in the rodent brain (14). We determined the presence of circRims2 by RT-qPCR in combination with ribonuclease R treatment in spinal cord tissue. Similar to what we observed from our RNA-seq analysis, we observed that Rims2 goes down significantly after injury (Figure 5). In an effort to find additional putative circRNAs altered following SCI, we annotated our data and found over 200 circRNA that were significantly modulated after SCI (see Table2). To better understand the potential role that these circRNAs play in axonal growth, we modulated the level of ADAR1, that inhibits the formation of circRNA. Using siRNA to block ADAR1 expression we show that cortical neurons grow more abundant and longer axons (Figure 6). This finding directly suggests that up-regulating the formation of circRNA leads to longer axonal outgrowth and suggests reduction of these circRNAs following SCI impairs axon regeneration. Conversely, the knockdown of ADAR1 may increase levels of circRNA in vivo and could promote axonal outgrowth. Hence, it is imperative to be able to delineate the function of a circRNA in the axons to help unravel its role in regulating axonal translation. We also used RNAscope to confirm the expression of circRims2 RNA in spinal cord tissue. In combination with RbnR treatment that digests linear mRNA, such as GAP-43, but retains circRNA, we found that Rims2 expression was preserved in the tissue (Figure 7). While little is known about the role of circRNAs in neuronal function, both miRNA and RNA-binding protein (RBP) binding functions have been described for circRNA. In addition, there are several other noted functions, including being transcriptional regulators and binding to ribosomes directly.

To identify potential mechanisms underlying circRNA on axon growth, we used bioinformatic analyses to predict which miR and RBPs could be binding to circRims2. We found multiple likely miRNAs that could bind to circRims2 that have also been reported to either target known mRNAs that modulates axonal regeneration or has been shown to mediate SCI-injury (see Table 3). Circular RNA interactome predict the miRNAs that can bind to circRims2 (62). Of those predicted miRNAs two can also bind to the 3’UTR of GAP-43, an mRNA known to be transported into axons for local translation. They are miR-192 and -377, each having 8 or 7 predicted binding sites, respectively, to circRims2. Axonally translated GAP-43 supports axonal elongation (9) and we detect GAP-43 mRNA in the axons by RNAscope.

**Table 3.** Predicted miR that can bind to circRims2. Using Circular RNA Interactome, we predicted which miR can bind to circRims2 and the number of binding sites. We present miR that are known to mediate SCI injury or whose targets are a known protein that is an effector for axonal regeneration.

| miRNA predictions | #binding sites | Mediated effects | Targets |
| --- | --- | --- | --- |
| miR-495 | 31 |  | STAT3 |
| miR-494 | 21 |  | Nogo |
| miR-142-5p | 13 | detected in sera of patients with SCI |  |
| miR-145 | 12 | negative regulator of astrogliosis |  |
| miR-21 | 10 | Regulate Axon Growth | BDNF;PTEN |
| miR-431 | 10 | Enhances axonal regeneration | Wnt signalling |
| miR-155 | 8 | Deletion promotes longer neurites | CREB1 |
| miR-331-3p | 8 | attenuates neuropathic pain | RAP1A |
| miR-192 | 8 |  | GAP-43 |
| miR-377 | 7 |  | GAP-43 |

Yayon et al utilize a method called Patch-Seq analysis which is single-cell RNA-seq from fixed tissues using iDISCO and advanced microscopy (NuNeX) to help spatially select VChIs neurons and detect changes in dendritic regions of these neurons after whisker deprivation model (68). By doing so, they have detected dendrite localized transcripts such as Elmo1, Msi2 and Tubb3 (β-III tubulin) which are down-regulated, while Arpp21 and Sema5b are up-regulated. From our findings, we observed that Tubb3 and Sema5b, which are detected in both dendrites and axons, go down significantly after injury. We detect no significant changes in Elmo1 or Msi2.

Intriguingly, Arpp21 goes down significantly after injury in our RNA-seq, by 4-fold and it is a putative circRNA. Further analyses are required to have a better understanding the consequences of these changes in potentially dendritic and axonal signatures. Similar to other reports such as Hanan et al. looking at circSLC8A1 in a Parkinson’s Disease model, we will follow-up our findings of circRims2 to elucidate it’s molecular mechanism to potentially modulate axonal outgrowth and to determine the miRNA and/or RBPs that it could be binding to for regulation of this outgrowth (69).

These observations predict a RNA-based regulatory pathway from circRNA Rims2→ miRNA (miR-192 or -377)→GAP-43 mRNA to control axonal levels of GAP-43 protein and regulate axonal outgrowth (Figure 8). The decreased levels of circRims2 after SCI indicate that there may local internal RNA-mechanisms that control axonal regeneration and RNA based therapies could play a role in neurorepair.

**Figure 8.**
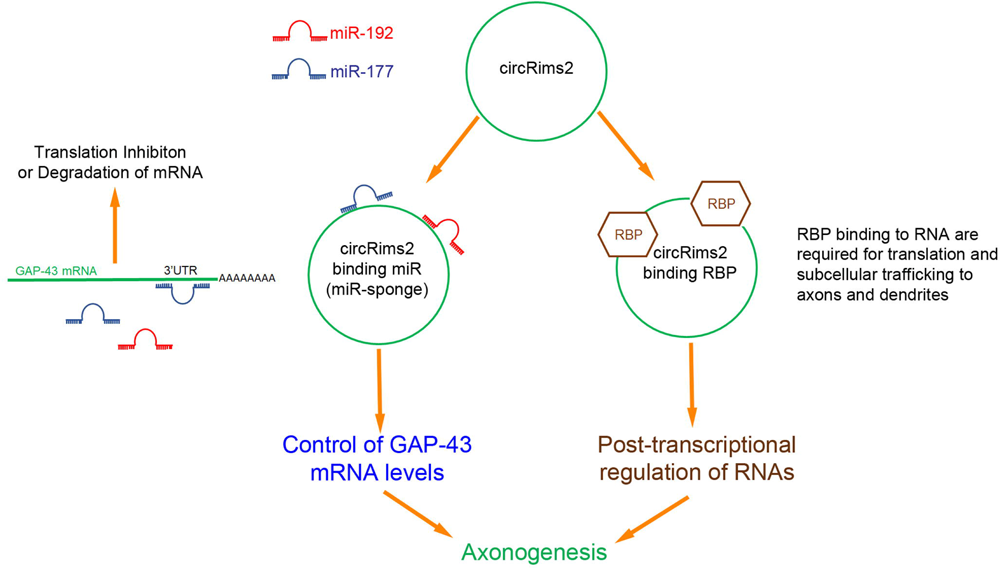
Inferred RNA pathway for circRims2 in modulating axonogenesis. **A.** By data mining, we identified 18 miR that could bind to circRims2 including miR-192 (red) and miR-177 (blue) that can target the 3’UTR of GAP-43 mRNA. mRNA bound to miR are either inhibited from being translated or degraded. **B.** circRNAs have been described to be miR-sponges. circRims2 could competitively bind miR-192 and -177 and prevent them from binding to GAP-43 mRNA, controlling expression of GAP-43 and promoting axonal outgrowth. **C.** circRims2 could also bind to RBPs, such as FUS that are known to mediate axonal outgrowth. RBPs can be involved in subcellular trafficking of RNA to axons and dendrites. RBPs are known to also regulate translation, hence another putative mechanism for regulating axonogenesis.

## Acknowledgements

The authors would like to thank Dr. Soheila Karimi at the University of Manitoba, Winnipeg, Canada for assistance with oligodendrocyte markers and Dr. Stella Tsirka at Stony Brook University, USA for assistance with microglial cell markers. We also thank Bindu Sundaresan, Connie Zhang and Jacqueline Giliberti at ACD for their assistance with RNA-scope detection of circRims2. Supported by NIH grant GM 137056 and NYS Spinal Cord Injury Research Program Contract number #C34460GG.

## Supplementary Figures

**Supplementary Figure 1 – Up-and down-regulated biological processes by GO analysis.**

From our RNA-seq we used GO analysis to determine biological processes that are being regulated. In the top figure in gold are pathways that are significantly up-regulated when comparing control to SCI. The lower figure in blue are pathways that are significantly down-regulated. The red arrows are pointing to pathways known to be involved with neurons.

**Supplementary Figure 2 - siRNA against ADAR1 compared to scrambled controls promote longer axons in rat primary cortical neurons-**

Graph from Figure 6C is replotted to show the distribution of points. This graph is the average of four independent experiments with standard deviations. Statistics is t-test (*p<0.05 and **p<0.01) comparing Control to ADAR siRNA (blue) at a fixed distance in the microgrooves. At 800 and 900 microns, ADAR siRNA (blue) were significantly longer compared to controls.

**Supplementary Figure 3 – Detection of ribosomes in the axons and growth cone tips of rat cortical neurons.**

We confirm expression of ribosomes in our cortical neurons by immunohistochemistry using an antibody against S6 Ribosomal Protein (green). We use Actin-Red to visualize the growth cones (white arrow). Dapi (blue) is staining the cell bodies and β-III tubulin immunohistochemistry to see the axons is in white.

**Supplementary Figure 4 – Prediction of 7 RBPs binding to circRims2.**

Using Circular RNA Interactome we predicted 7 RBPs that could bind to circRims2. The number in each column represents the number of binding sites. Fused in Sarcoma (FUS) has 16 binding sites on circRims2.

