## SupplementaryFigures for "Spinal Cord Injury regulates circular RNA expression in axons"

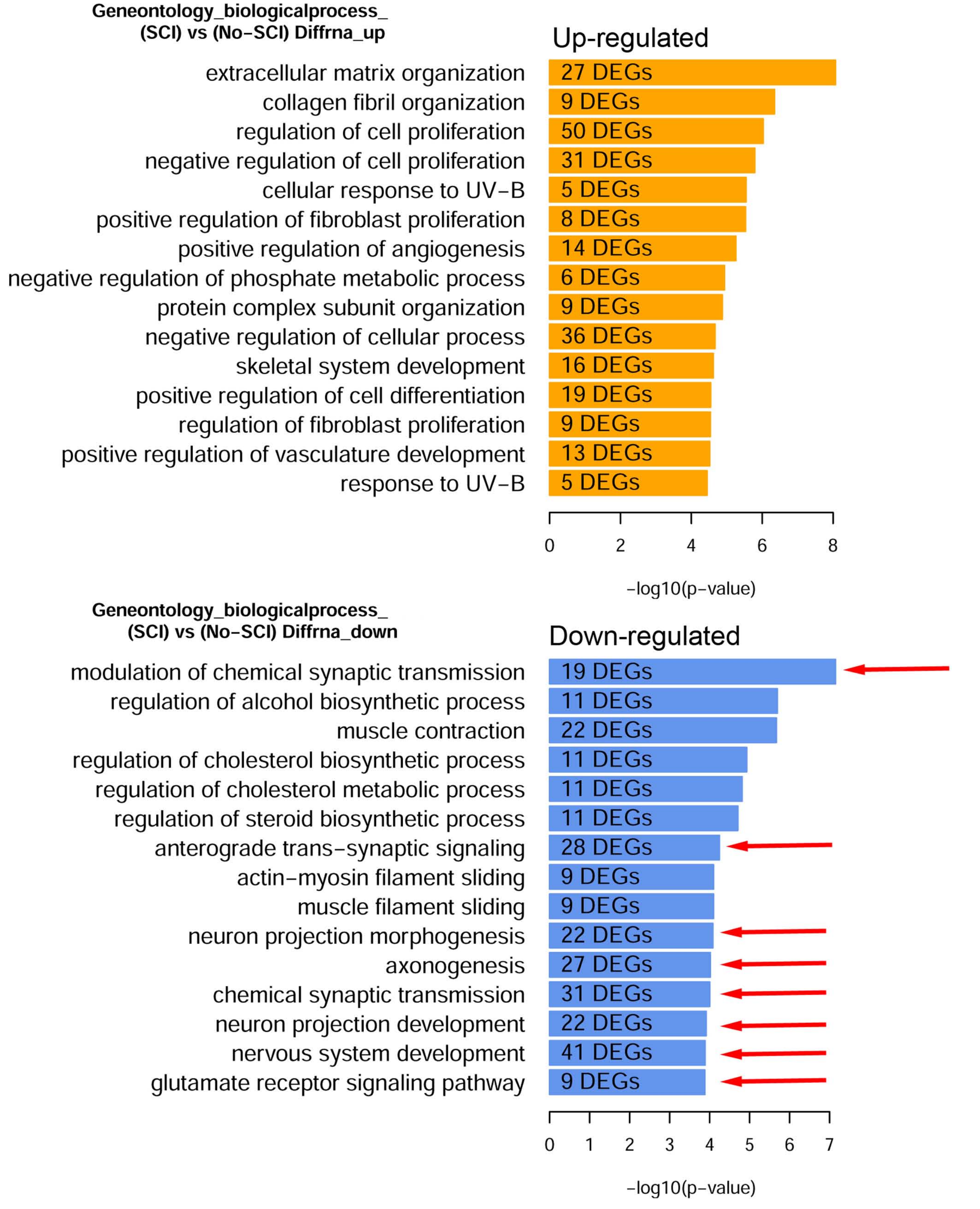
Geneontology_biologicalprocess_
(SCI) vs (No-SCI) Diffrna_up
Up-regulated
extracellular matrix organization
collagen fibril organization
regulation of cell proliferation
negative regulation of cell proliferation
cellular response to UV-B
positive regulation of fibroblast proliferation
positive regulation of angiogenesis
negative regulation of phosphate metabolic process
protein complex subunit organization
negative regulation of cellular process
skeletal system development
positive regulation of cell differentiation
regulation of fibroblast proliferation
positive regulation of vasculature development
response to UV-B
27 DEGs
9 DEGs
50 DEGs
31 DEGs
5 DEGs
8 DEGs
14 DEGs
6 DEGs
9 DEGs
36 DEGs
16 DEGs
19 DEGs
9 DEGs
13 DEGs
5 DEGs
0
2
4
6
8
-log10(p-value)
Geneontology_biologicalprocess_
(SCI) vs (No-SCI) Diffrna_down
Down-regulated
modulation of chemical synaptic transmission
regulation of alcohol biosynthetic process
muscle contraction
regulation of cholesterol biosynthetic process
regulation of cholesterol metabolic process
regulation of steroid biosynthetic process
anterograde trans-synaptic signaling
actin-myosin filament sliding
muscle filament sliding
neuron projection morphogenesis
axonogenesis
chemical synaptic transmission
neuron projection development
nervous system development
glutamate receptor signaling pathway
19 DEGs
11 DEGs
22 DEGs
11 DEGs
11 DEGs
11 DEGs
28 DEGs
9 DEGs
9 DEGs
22 DEGs
27 DEGs
31 DEGs
22 DEGs
41 DEGs
9 DEGs
0
1
2
3
4
5
6
7
-log10(p-value)

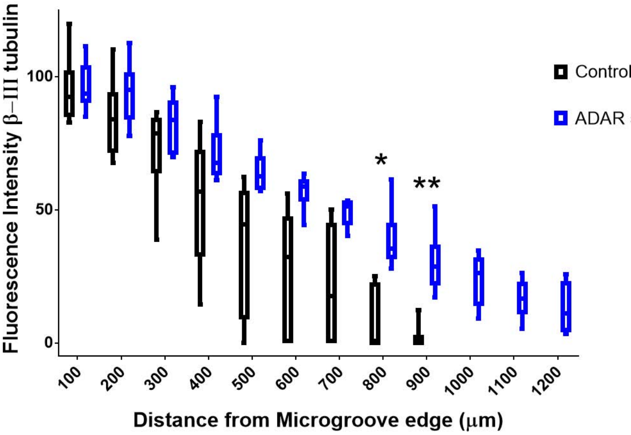

Fluorescence Intensity β–III tubulin
100
50
0
*
**
Control
ADAR
100
200
300
400
500
600
700
800
900
1000
1100
1200
Distance from Microgroove edge (μm)

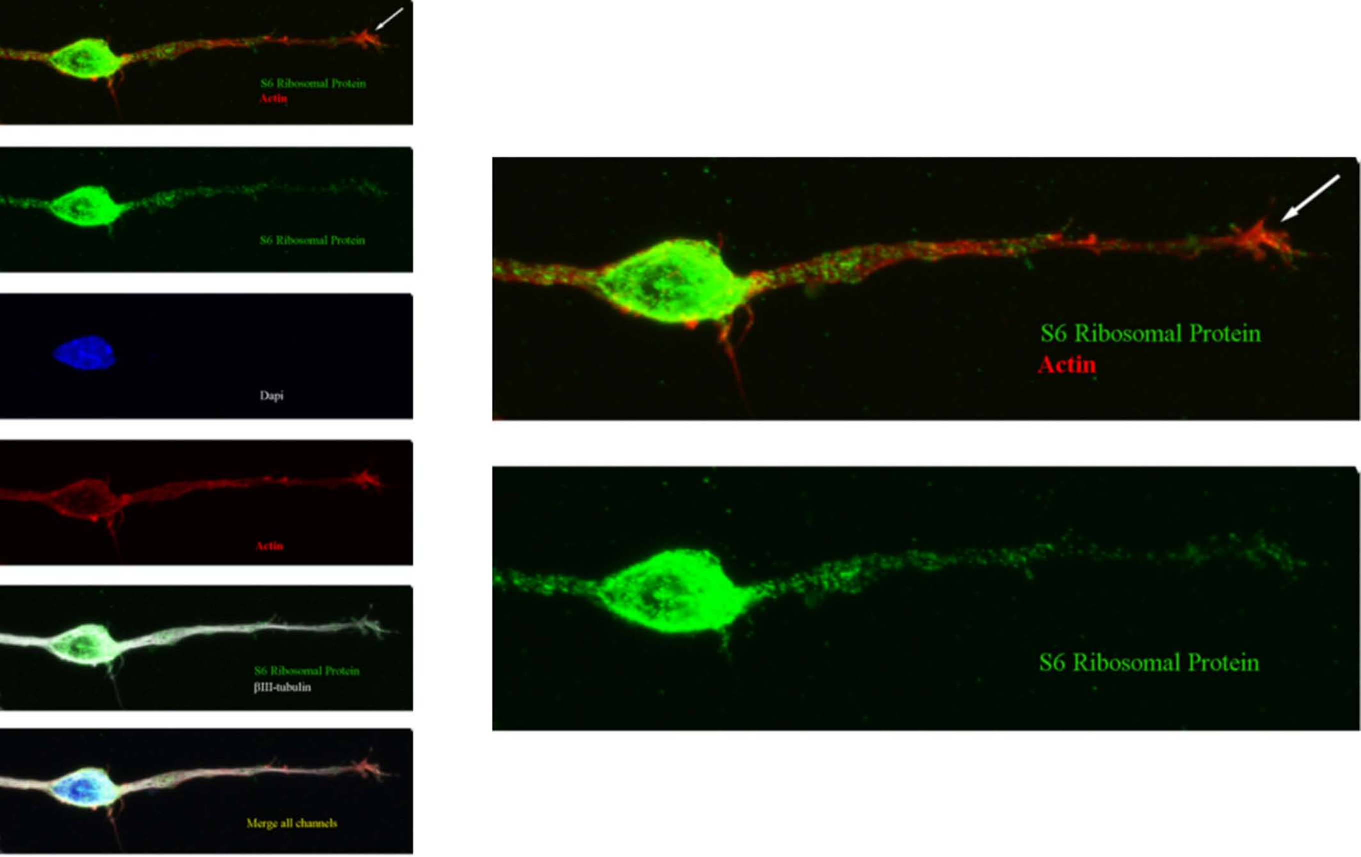
S6 Ribosomal Protein
Actin
S6 Ribosomal Protein
Dapi
Actin
S6 Ribosomal Protein
βIII-tubulin
Merge all channels
S6 Ribosomal Protein
Actin
S6 Ribosomal Protein

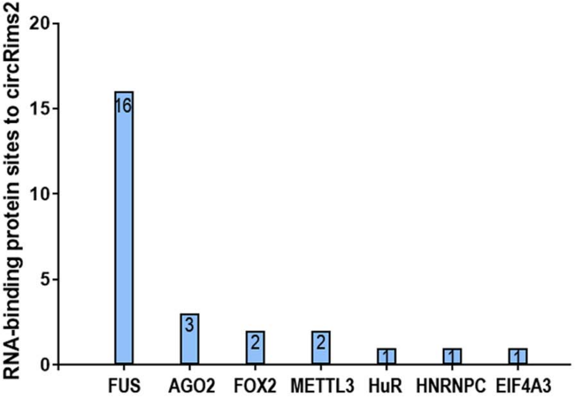

RNA-binding protein sites to circRims2
20
15
10
5
0
16
3
2
2
1
1
1
FUS
AGO2
FOX2
METTL3
HuR
HNRNPC
EIF4A3

**Supplementary Table 1 – Additional markers for spinal interneurons and motor neurons.** These are additional markers for spinal interneurons and motor neurons that have been reported. They are detected in low abundance. Intriguingly, ESRRG which was significantly down-regulated after injury is also a putative circRNA. Adjusted P value and fold change are given.

| DEG | No-SCI | SCI | P-Adj | Log2 Fold |
|---|---|---|---|---|
| SLC17A6 | 2.14437 | 1.669477 | 0.419391 | -0.3611577 |
| SLC32A1 | 1.800888 | 1.191043 | 0.140346 | -0.596483 |
| ESRRG | 7.147771 | 1.785862 | ***0.000329625*** | -2.000873 |
| EBF1 | 38.69875 | 22.71682 | 0.0659941 | -0.7685261 |
| TAC2 | **Not Detected** | | | |
| MAF | 3.63676 | 8.09915 | 0.1915 | 1.155117 |
| PKD2 | 2.85083 | 2.93693 | 0.959094 | 0.04292679 |
| CHAT | **Not Detected** | | | |
| PDK1I2 | **Not Detected** | | | |
| NTS | 3.3992 | 1.533663 | ***0.014*** | -1.148214 |
| CCK | 11.79828 | 17.46138 | 0.576153 | 0.565591 |
| EN1 | **Not Detected** | | | |
| VSX2 | **Not Detected** | | | |
| HOXA10 | 5.10762 | 2.84342 | 0.158656 | -0.8450239 |
